# Mechanistic Insights into Crossover-Dependent Stability and Exceptional Resistance of PX/JX Nanostructures to DNase I using Enhanced Sampling

**DOI:** 10.1101/2025.04.17.649409

**Authors:** Sandip Mandal, Arun Richard Chandrasekaran, Prabal K. Maiti

## Abstract

Designing biostable DNA nanocarriers for precise therapeutic delivery remains a key challenge in DNA nanotechnology due to susceptibility to nuclease degradation. Multi-stranded DNA nanostructures, such as Paranemic crossover (PX) DNA, show enhanced biostability compared to native duplex DNA due to their unique topology. However, the molecular origin of their exceptional nuclease resistance is still unknown. Using atomistic MD simulation and enhanced sampling, we uncover the molecular origins of PX-DNA’s superior resistance over JX-DNA and dsDNA. Our findings reveal that PX-DNA’s six crossover points induce an overtwisted helix and narrower minor groove, leading to reduced DNase I binding affinity (∼ +5 kcal/mol for PX-DNA vs. ∼ −17 kcal/mol for dsDNA). Mechanical properties, such as the stretch modulus (*γ_G_*), confirm enhanced structural rigidity of PX-DNA (4805 pN), while residence time calculations using *τ*-RAMD further highlight shorter residence time of DNase I nuclease on PX-DNA. This is the first theoretical study to explore crossover-dependent biostability, mechanical properties, and residence time of PX/JX nanostructures in the presence of a nuclease using enhanced sampling. Our findings highlight that strategically positioned crossover points can regulate DNA stability against nuclease degradation at the nanoscale and lead to better design of biostable DNA nanostructures for medicinal applications.

## Introduction

In addition to playing a fundamental role in biological heredity, DNA is the most popular molecular building block for self-assembly and nanofabrication and enables the construction of diverse 1D, 2D, and 3D DNA nanostructures.^1^ Beyond biocompatibility, DNA has several advantages, including a clearly defined geometry, predictable and programmable Watson-Crick (WC) base-pair complementarity, diverse sequences, thermostability, and easily accessible chemical and enzymatic modification tools. ^2–4^ The success of mRNA vaccines during the COVID-19 pandemic resulted in a new era of therapeutic nucleic acids. These nucleic acid-based drugs enable precise, gene-specific personalized treatments for a variety of genetic diseases, revolutionizing modern medicine.^5^ However, their stability is still a significant challenge because the human body has evolved with multiple enzymes and nucleases to degrade unwanted DNA/RNA, limiting therapeutic efficacy.^6^ The susceptibility of the native dsDNA-based nanostructures to nucleases in the physiological environment is a major drawback for use in drug delivery. ^7–9^ To overcome the key challenges in the stability of nucleic acid-based drugs and nanostructures as drug delivery vehicles, various strategies have been developed to extend their lifespan, including encapsulation, thymine cross-linking, G-quadruplex formation, and synthetic nucleotide modifications.^10–13^ Even though some experimental studies have explored the nuclease resistance of DNA nanostructures, ^14–16^ their interactions with the nucleases still remain elusive. Chandrasekaran et al.,^17^ investigated the enzymatic degradation of native dsDNA (Figure 1A), DX, and PX-DNA (Figure 1B) motifs in the presence of DNase I nuclease. Their findings revealed that PX-DNA exhibited superior nuclease resistance, remaining intact for hours, while 90% of dsDNA degraded quickly within minutes. These findings suggest that DNA nanostructure topology influences binding affinity, with PX-DNA and JX(1,2,3,4) DNA potentially obstructing nuclease binding sites, prolonging their stability in physiological environments from minutes to hours.

**Figure 1:**
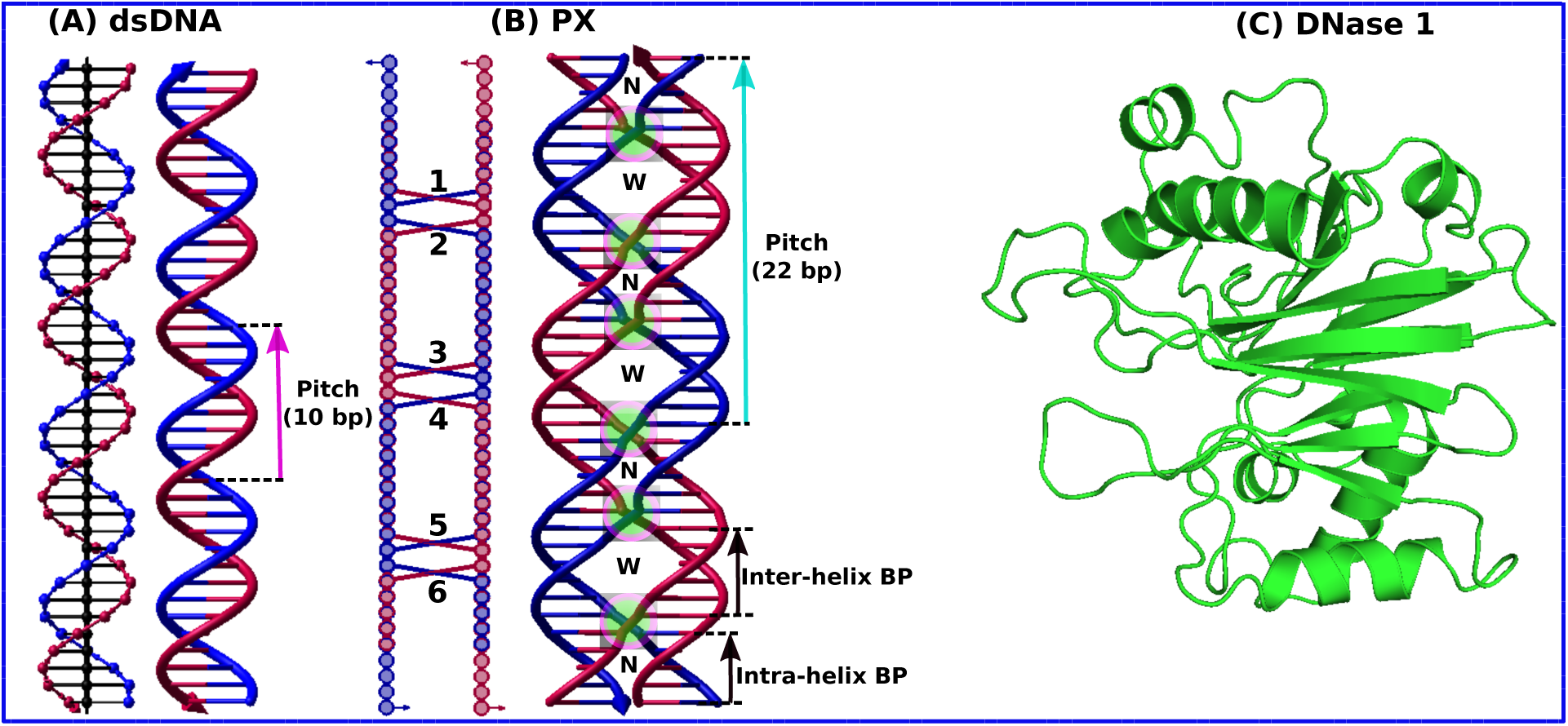
Schematics for (A) native dsDNA, (B) paranemic crossover (PX-DNA). The six crossover locations are shown as transparent green circles. The helical pitch of dsDNA is 10 base pairs, whereas for PX-DNA, its 22 base pairs. (C) Represents the initial pdb structure of the DNase I nuclease, which cleaves phosphodiester bonds between nucleotides.

DNA nanostructures are highly programmable, enabling exact control over shape, size, and versatile functionality. ^1,18^ Since Ned Seeman first proposed the concept of DNA nanotechnology 40 years ago, various DNA nanostructures,^19,20^ such as DNA origami nanotubes, flasks and cages with different sizes and shapes, have been self-assembled and employed in various applications including molecular biology, drug delivery, ^21–24^ gene therapy^25,26^ and biosensings.^27,28^ For drug delivery applications, DNA nanostructures (PX/JX) provide high intrinsic biocompatibility and non-cytotoxicity while protecting the drugs from enzymatic damage (such as by DNase I endonuclease that cleaves the phosphodiester bonds between nucleotides, as shown in Figure 1C),^18,29,30^ due to their topological uniqueness from the native dsDNA. To improve the biostability of DNA/RNA nanostructures, previous studies have also utilized polymers, protein coatings, modified nucleotides, nucleobase mimics like cyanuric acid, and cross-linking of component strands.^10,11,31–34^ Beyond these strategies, the possibility of designing new DNA nanostructures that can change their biostability just by tuning their topology has been largely overlooked.

Several multi-crossover DNA structures have been designed, including double crossover (DX), triplex crossover (TX), and multi-star motifs.^35^ A rarer motif, Paranemic crossover (PX) DNA, features a coaxial four-stranded structure with two connected double-helical domains, where every nucleotide forms Watson-Crick (WC) pairs. ^36,37^ PX-DNA forms through reciprocal exchange between strands of the same polarity at every potential position where two juxtaposed DNA duplexes are placed side by side. Each helix of the PX-DNA has an alternating wide major (W) and narrow minor (N) groove separation (shown in Figure 1B). Previously researchers have engineered PX-DNA with various major:minor groove separations (W:N), with the most stable complexes containing 6, 7, or 8 nucleotides in the major groove and 5 nucleotides in the minor groove (PX 6:5, 7:5, and 8:5). ^36,38–40^ PX 6:5, one of the stable variants that has been widely used in the field of structural DNA nanotechnology,^41–47^ has an 11 base pair half-pitch (22 base-pair per full-pitch), closely matching with the 10.5 base pairs per turn of B-form dsDNA.^44,45,48^ Six crossover sites in the PX 6:5 molecule lead to the structure depicted in Figure 2A (upper panel) (For simplicity, PX 6:5 is referred to as PX-DNA throughout this article). JX1, a topoisomer of PX-DNA, forms by removing its central crossover from the PX-DNA. The JXn structure is related to PX by containing n adjacent sites where the backbones of the two parallel double helices juxtapose without crossing over. Thus, JX2, JX3, and JX4 form by eliminating two, three, or four consecutive crossovers as shown in Figure 2(B, C, D, and E) (upper panel). In DNA nanotechnology, PX-DNA motifs have been used to construct triangles, octahedra, 1D and 2D arrays,^45,49–54^ as well as for molecular assembly and DNA computation in nanomechanical devices.^44,48^ Despite their broad appeal in nanotechnology and biology, PX-DNA’s atomistic-level biostability remains unexplored.

**Figure 2:**
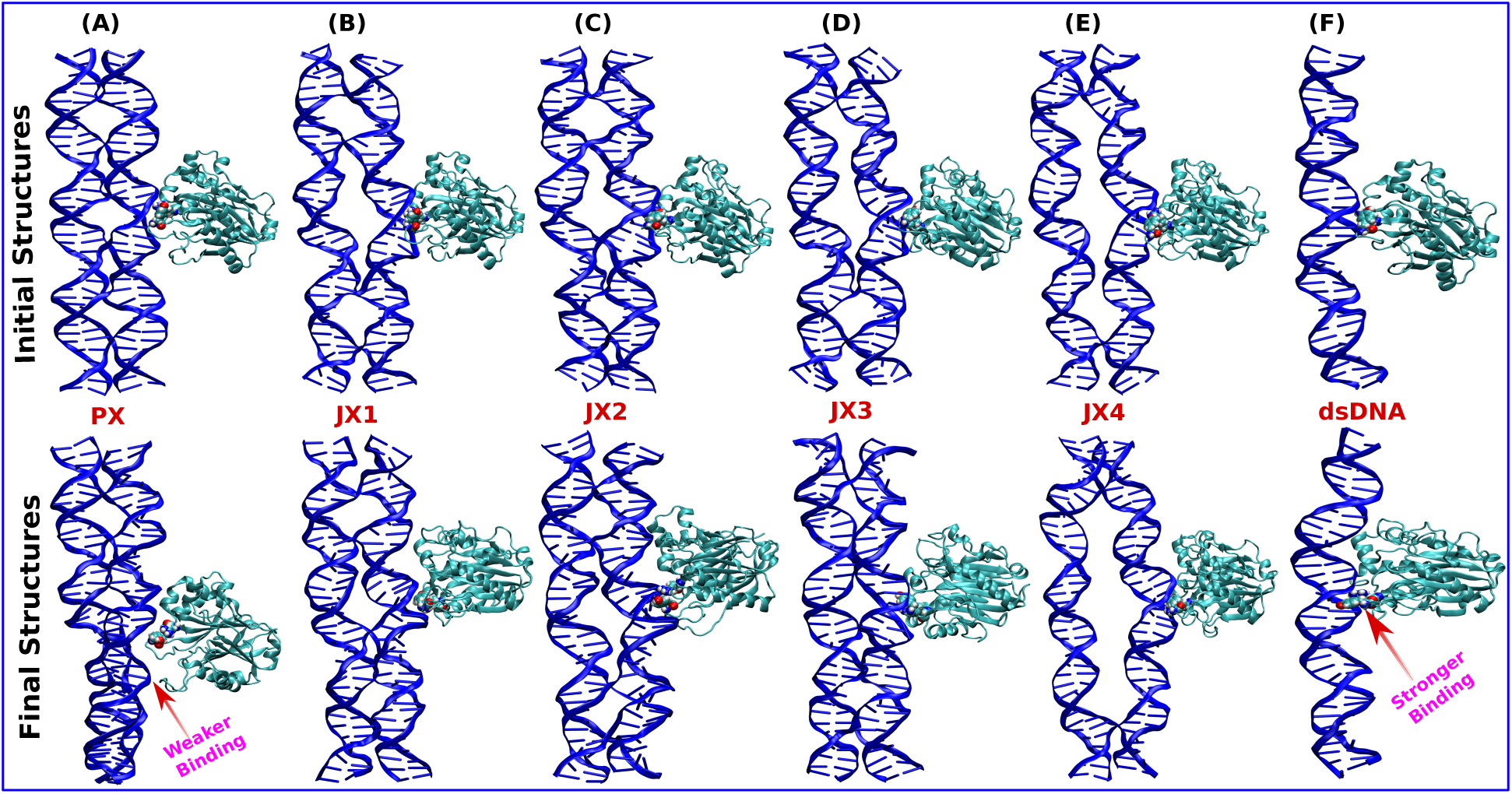
Instantaneous snapshots of the initial and final conformations of the six DNA-DNase I complexes from the 300 ns long MD simulations. The upper and lower panel corresponds to the initial and final structures of (A) PX-DNase I, (B) JX1-DNase I, (C) JX2-DNase I, (D) JX3-DNase I, (E) JX4-DNase I and (F) dsDNA-DNase I complexes. The final conformations in the lower panel suggest that dsDNA-DNase I has a stronger binding affinity in the dsDNA grooves, and PX-DNase I has the least binding affinity.

In this work, we investigated the molecular interactions between DNase I nuclease and DNA nanostructures (PX/JX’s(1,2,3,4) and native dsDNA) using a 300 ns long atomistic MD simulation trajectory (Figure 2, Lower panel). The trajectory analysis conveys that PX-DNA has significantly higher nuclease resistance, driven by its maximum number of crossover points and greater rigidity modulus. Each additional crossover enhanced protection against nuclease degradation, highlighting the potential to engineer highly stable DNA nanostructures using PX-DNA.

## Results and Discussion

### Structural Properties

We assessed the stability of native dsDNA and PX/JX-DNA nanostructures upon DNase I nuclease binding by calculating RMSD over 300 ns of unbiased MD simulations, using their energy-minimized initial structures as references as shown in Figure 3A. The RMSD of a structure at time t, relative to its reference structure, is 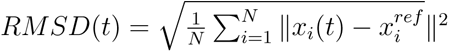, where the coordinates of the N atoms in the structure are denoted by {*x_i_*(*t*)}, and the corresponding coordinates of the initial energy-minimized structure are denoted by 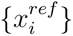. PX-DNA, with six crossover points, exhibits higher RMSD due to over-twisting caused by strong electrostatic repulsion and rigidity, despite being the most stable structure in the absence of DNase I.^39^ In contrast, dsDNA shows the highest structural fluctuations due to its flexibility and strong binding affinity to the DNase I, leading to bending. The JX-DNA structures, with fewer crossovers, show intermediate RMSD values, reflecting reduced rigidity with decreasing crossover points. The averaged RMSD values are displayed in a histogram plot for better quantitative understanding (Figure 3B). All the measurements of various physical quantities in this manuscript are averages of the last 100 ns of the total 300 ns long MD simulation trajectories (as indicated by gray shaded regions in Figure 3A, C and E).

**Figure 3:**
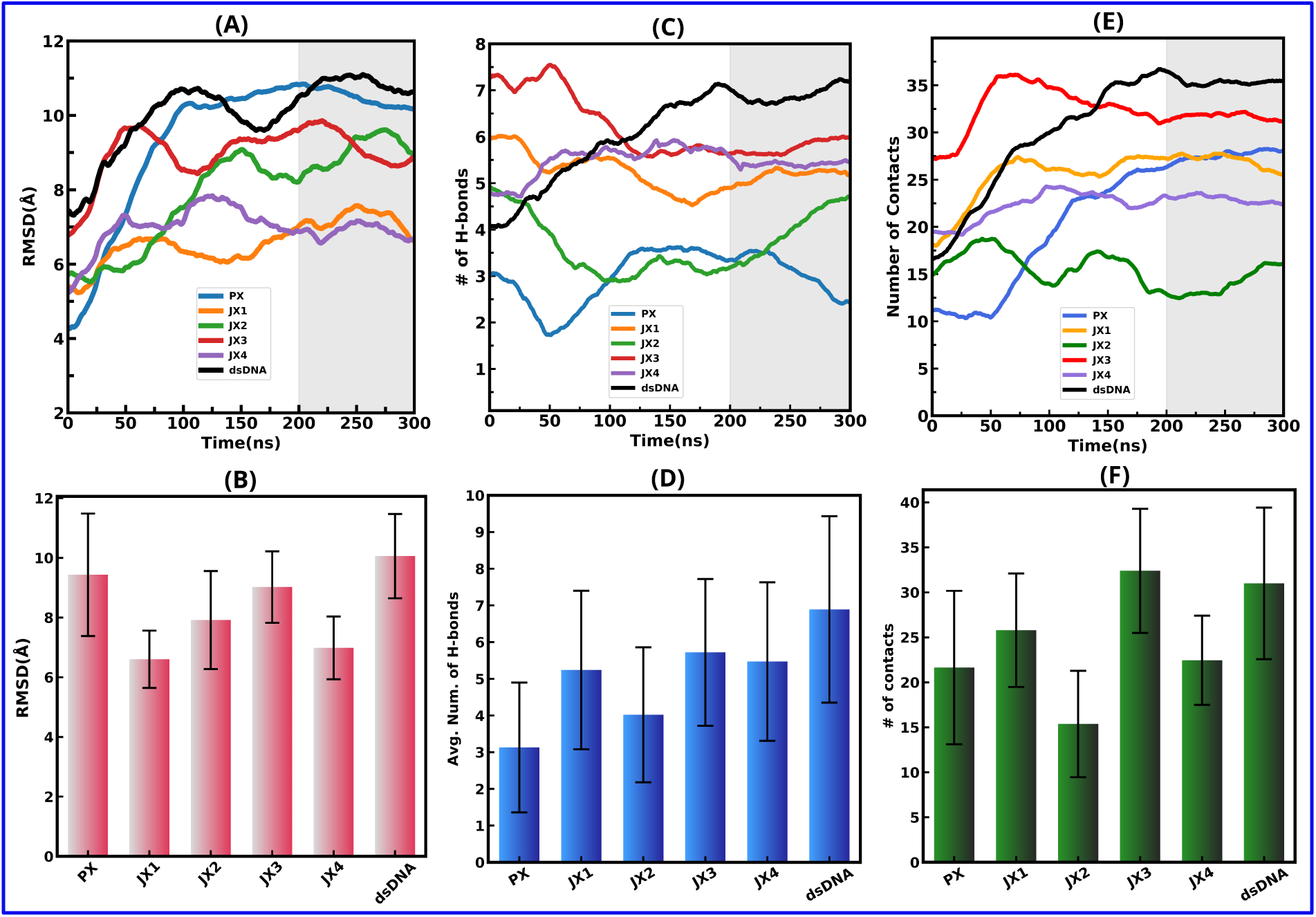
(A) RMSD of dsDNA and PX/JX-DNA complexes with DNase I over 300 ns MD simulations. (B) The average and standard deviation of the RMSD for quantitative comparison of the stability of all the six complexes. (C) Hydrogen bond (HBs) count between DNA nanostructures and DNase I. (D) The time average of the number of HBs count. (E) Time evolution of the number of contacts between the dsDNA, PX/JX-DNA, and DNase I complexes within the cutoff distance of 3.5 Å. (F) The time average of the number of contacts. Gray shading in (A), (C), and (E) highlights the final 100 ns of the 300 ns simulation.

Hydrogen bonds (HBs) play a critical role in DNase I nuclease binding to DNA nanostructures. Using CPPTRAJ^55^ in AMBER20,^56^ we calculated the average HBs (as listed in Table-1) and the number of contacts between DNase I nuclease residues and DNA nanostructures (PX/JX), from the last 100 ns of the total 300 ns MD simulations (with distance cutoff: 3.5 Å, angle cutoff: 120*^o^*, as recommended by IUPAC^57^). Native dsDNA shows the highest number of HBs, indicating deeper minor groove penetration and stronger binding affinity as compared to JX’s and PX-DNA (Figure 3C-D). The PX-DNA exhibits the fewest HBs and contacts, consistent with its weaker binding affinity and enhanced rigidity. The JX topoisomers show an intermediate number of contacts, with JX3 displaying contacts similar to dsDNA due to increased flexibility with reduced crossovers. These findings underscore DNase I’s preference for dsDNA over crossover-rich PX-DNA motifs.

**Table 1:**
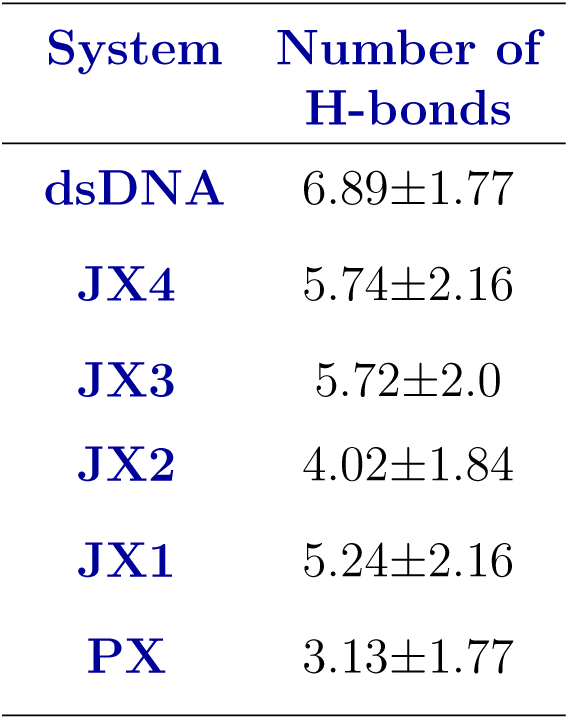
Average number of hydrogen bonds at 300 K using the last 100 ns from the 300 ns long MD simulation trajectory for all the six DNA nanostructure-DNase I complexes.

We analyzed DNase I contacts with all six DNA nanostructures (PX, JX, and dsDNA) using the CPPTRAJ tool provided with AMBER20 software, taking a distance cut-off of 3.5 Å (Figure 3E). The number of contacts shows a similar trend as the number of HBs, with DNase I forming 19.54 ± 2.56 contacts with dsDNA and 11.60 ± 3.17 with PX-DNA (Figure 3F). These results indicate DNase I’s preference for dsDNA motifs for binding (Figure 4A-B).

**Figure 4:**
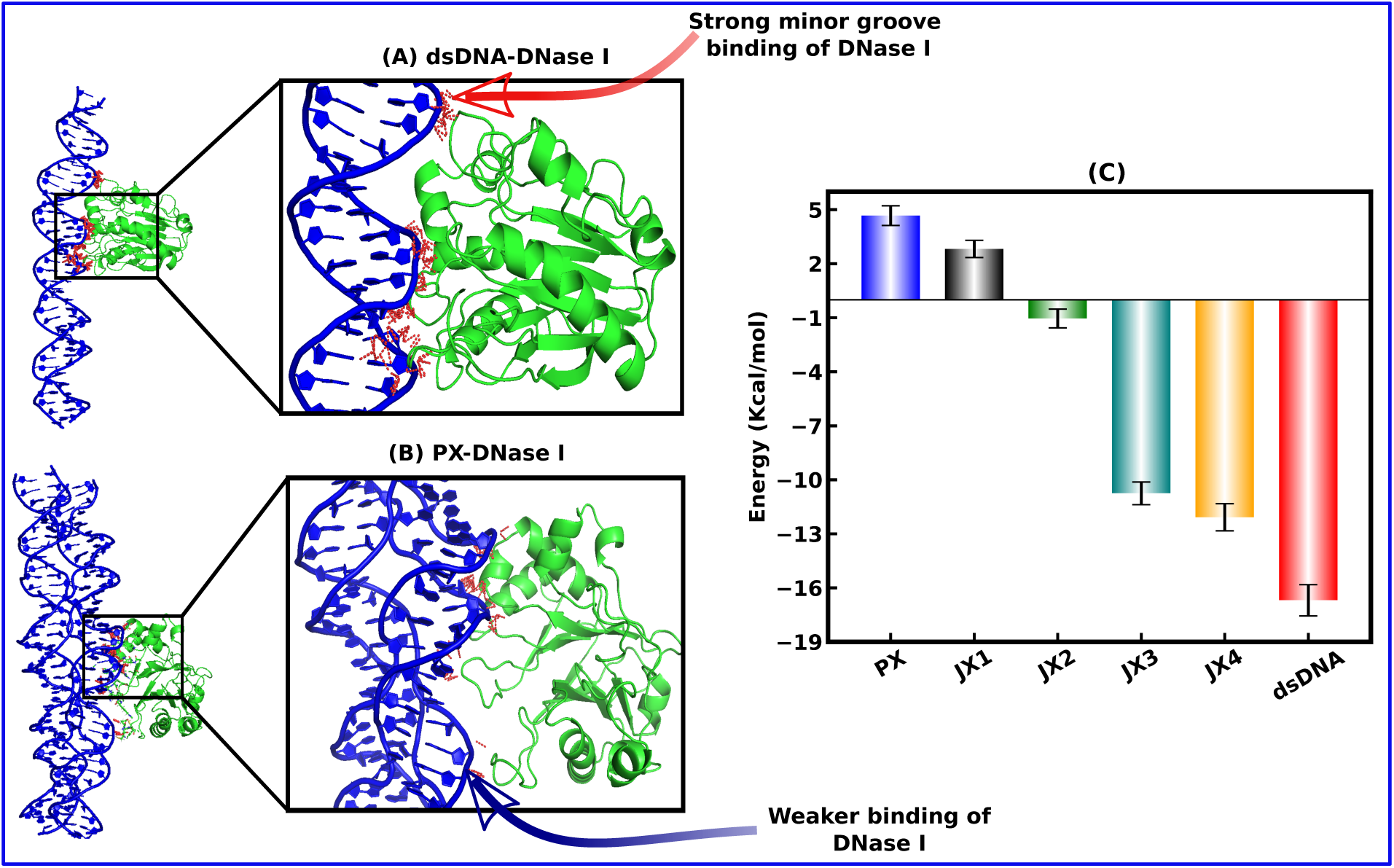
(A)Snapshots for binding of DNase I to dsDNA grooves through hydrogen bonding. (B) Snapshots for DNase I binding to PX-DNA with weaker binding affinity and fewer hydrogen bonds compared to dsDNA. The red dashed lines indicate the hydrogen bonds. (C) Total binding free energy from MMGBSA analysis for all six DNA-DNase I complexes using the last 50 ns from the 300 ns long MD trajectory, in kcal/mol units.

## Energetic Properties

### Binding Energy Calculation using MMGBSA

MMGBSA (Molecular Mechanics-Generalized Born Surface Area) provides a useful approach to estimating the energetics of ligand and receptor bindings. The obtained binding free energy provides deep insight into the binding affinity and stability. The binding free energies of all the six DNA-DNase I complexes were calculated using the MMGBSA ^58–62^ approach with the MMPBSA.py^63^ module of AMBER20.^56^ The binding free energy can be calculated as:

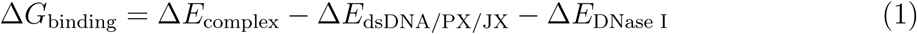

where Δ*E*_complex_, Δ*E*_dsDNA/PX/JX_ and Δ*E*_DNase I_ represents the free energies of all six DNA-DNase I complexes, and dsDNA/PX/JX and DNase I individual receptors.

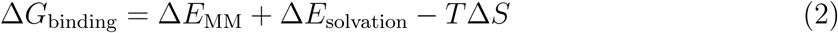

Here, Δ*E*_MM_ refers to the change in gas-phase molecular mechanics energy, *T* Δ*S* is the entropy change, and Δ*E*_solvation_ is the change in solvation energy after ligand binding. Δ*E*_MM_ can be further expressed as:

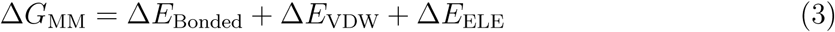

Understanding the binding affinity of DNA nanostructures is key to explaining their degradation by DNase I. Prior studies by Chandrasekaran et al.^17^ showed DNase I readily degrades dsDNA within a few minutes, but multi-crossover PX/JX-DNA are stable for up to hours. Therefore, using the MMGBSA approach, we found DNase I binds to dsDNA with stronger affinity (∼-17 kcal/mol), approximately 22 kcal/mol higher than PX-DNA (∼+5 kcal/mol) (Figure 4C). This reflects the remarkable nuclease resistance of PX-DNA, driven by its crossover-rich unique topology and helical parameters, compared to dsDNA and JX’s (JX1-JX4). To evaluate the DNase I’s binding affinity to the native dsDNA and DNA nanostructures (PX and JX’s), we estimated the binding energy of the complexes using equation-1, 2 and 3 and the observed values are listed in Table-2.

**Table 2:** Different energy components of binding free energy from MMGBSA analysis at 300 K using the last 50 ns from the 300 ns long MD simulation trajectory in kcal/mol units.

| System | $\Delta E_{\text{VDW}}$ | $\Delta E_{\text{ELE}}$ | $\Delta E_{\text{GB}}$ | $\Delta G_{\text{Binding Energy}}$ |
| --- | --- | --- | --- | --- |
| **PX** | $-59.65 \pm 4.62$ | $10841.81 \pm 224.89$ | $-10769.84 \pm 223.04$ | $4.67 \pm 0.35$ |
| **JX1** | $-66.80 \pm 6.72$ | $10390.60 \pm 271.53$ | $-10312.79 \pm 267.27$ | $2.82 \pm 0.18$ |
| **JX2** | $-50.50 \pm 4.29$ | $10111.04 \pm 190.46$ | $-10055.55 \pm 187.99$ | $-1.04 \pm 0.22$ |
| **JX3** | $-63.53 \pm 4.87$ | $10222.61 \pm 161.28$ | $-10162.64 \pm 159.40$ | $-10.75 \pm 0.28$ |
| **JX4** | $-61.84 \pm 5.47$ | $9869.37 \pm 225.14$ | $-9811.62 \pm 224.41$ | $-12.08 \pm 0.23$ |
| **dsDNA** | $-74.77 \pm 5.11$ | $6006.60 \pm 96.52$ | $-5938.11 \pm 93.83$ | $-16.69 \pm 0.27$ |

The electrostatic energy (EEL) between PX-DNA and DNase I is strongly repulsive because DNase I is a negatively charged molecule, and the PX-DNA motif carries higher negative charges (the distance between the two double helices is the shortest, and the presence of crossovers brings the negative charge centers closer), and has a unique topology with six crossover points. Native dsDNA, in contrast, shows stronger van der Waals (VDW) interactions, indicating closer binding in its minor groove (Figure S1A-B). PX-DNA also exhibits higher Generalized-Born solvation energy (EGB), partially countering electrostatic repulsion (Figure S1C). At the same time, the native dsDNA-DNase I complex has decreased its solvation energy by establishing stronger hydrogen bonds, as seen by the higher negative non-polar solvation-free energy (Figure S1D).

Overall, based on the various non-bonded interaction energies from the unbiased simulation, MMGBSA calculations confirm DNase I’s strong affinity for native dsDNA and its inability to degrade PX-DNA, aligning with previous experimental findings.^17^ The JX-DNA motifs show intermediate binding affinity, highlighting a crossover-dependent trend in nuclease resistance. Therefore, for further enhanced sampling calculations (JE-SMD and US), we have considered two extreme systems, dsDNA and PX-DNA, with the highest and lowest binding affinities, as suggested by MMGBSA results.

### Binding Free Energy Calculation using Jarzynski Equality

Steered Molecular Dynamics (SMD), quantifies DNase I binding affinity to DNA nanostructures by measuring the non-equilibrium pulling work. Consider a thermally stable system in contact with a heat reservoir at a given temperature T. Let *λ* be a controllable macroscopic parameter(reaction coordinate) that alters the system’s state. If a system is driven isothermally from an initial equilibrium state A (consistent with *λ* = *λ_i_*) to B (consistent with *λ* = *λ_f_*) following a particular protocol *λ*(t): *λ_i_*→ *λ_f_*, the work performed is:

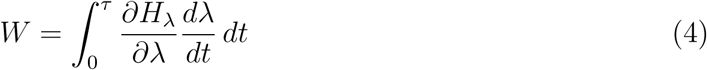

Here, *τ* denotes the transition time, and *H_λ_* represents how the system’s energy changes with the parameter *λ*. An ensemble of work values (W) can be obtained by repeating the above process multiple times. Jarzynski first developed an equality connecting the nonequilibrium work performed using SMD and the equilibrium free energy difference (ΔF) between states A and B as given by:

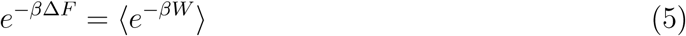

where ⟨·⟩ denotes the ensemble average over the work (*W*) values and 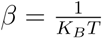. In this case, the ensemble average is calculated over multiple executions of the system’s non-equilibrium work performed during pulling simulations. The work was computed for each pulling process as the integral of the force applied over the pulling distance to the target molecule’s center-of-mass (COM) as:

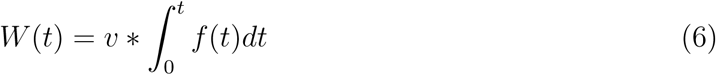

To solve computing issues caused by small work values in the exponential term, we use a cumulant expansion based on a Gaussian distribution as an approximation. Using Jarzynski’s equality, we can determine the free energy profile along the reaction coordinate (*λ*). When the DNase I achieves quasistatic equilibrium along this coordinate, all other degrees of freedom become entirely relaxed. SMD simulations are used to sample the reaction pathway and derive the Potential of Mean Force (PMF) with the stiff-spring assumption as:

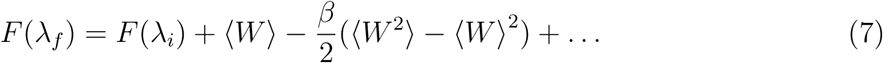

Therefore, Jarzynski’s Equality (JE) provides a theoretical framework for calculating the binding free energy difference by exponentially averaging the non-equilibrium works performed between bound and unbound states using Steered Molecular Dynamics (SMD). Therefore, JE computes the potential of mean force (PMF) by pulling the DNase I enzyme away from the binding regions of the PX-DNA and dsDNA. The non-equilibrium work (equation-4,5,6) was performed over 10 trajectories at very slow pulling speeds (0.1 nm/ns and 1 nm/ns) with a spring force constant of 3000 kcal/mol (Figure 5(A-F)). However, significant variations in the work that deviates from the Gaussian distribution lead to poorly converged PMFs using only the first two cumulants of the work distributions (equation-7). Thus, we used the exponential averaging form of Jarzynski’s Equality (provided by equation-5) to improve the accuracy. The PMF flattened beyond 4 nm, indicating the unbound state of DNase I. Using JE, binding free energies were calculated as Δ*G_dsDNA_* = +63.70 kcal/mol and Δ*G_PX−DNA_* = +27.05 kcal/mol, at 0.1 nm/ns, and Δ*G_dsDNA_* = +132.04 kcal/mol and Δ*G_PX−DNA_* = +75.20 kcal/mol at 1 nm/ns velocity, confirming the higher binding affinity of native dsDNA for DNase I as compared to PX-DNA.

**Figure 5:**
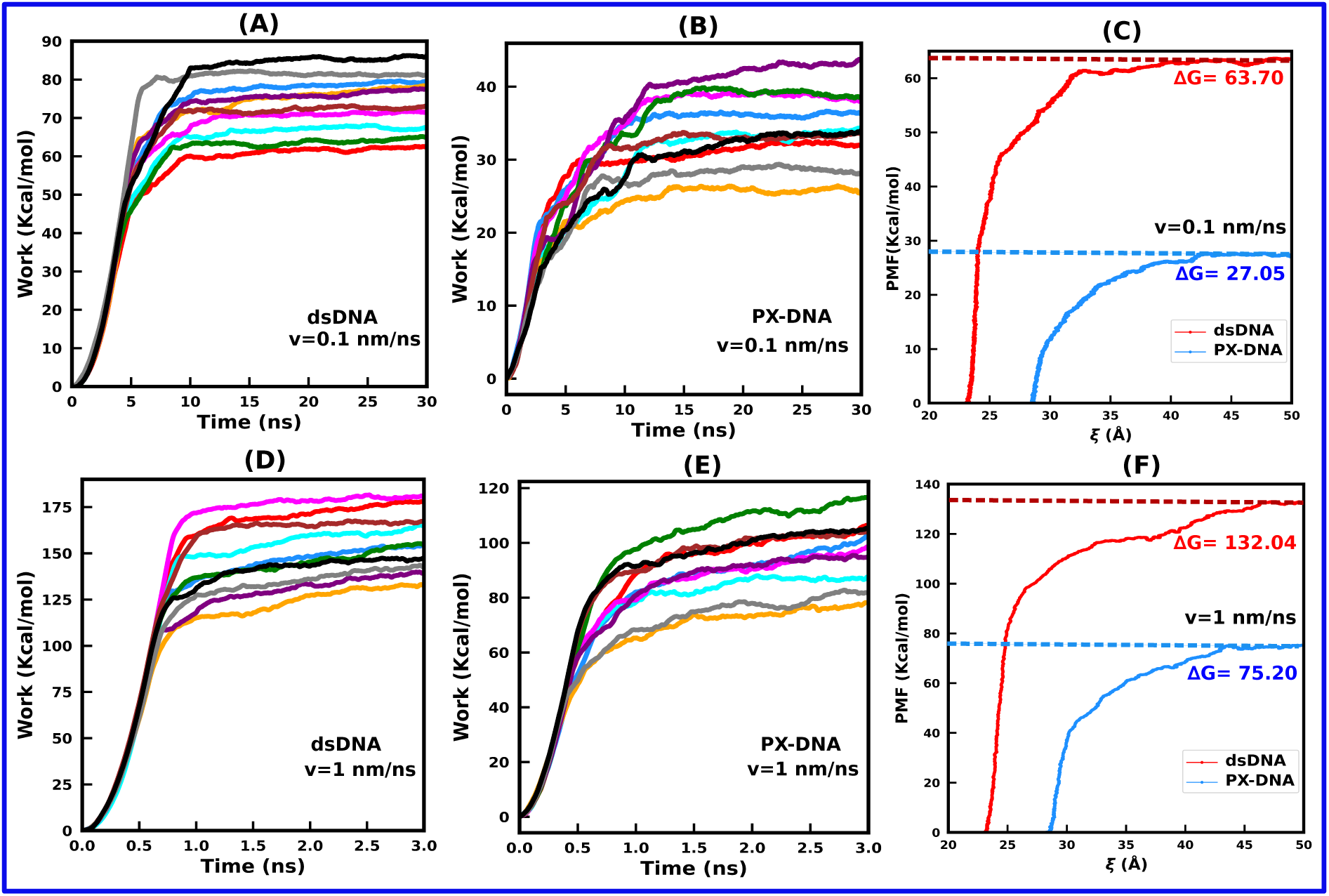
PMF and work profiles for slow pulling of the DNase I from the binding region of dsDNA and PX-DNA with a velocity of 0.1 nm/ns and 1 nm/ns by utilizing the SMD and JE for all 10 SMD trajectories. [A, B] the non-equilibrium work profiles of the DNase I dissociation from dsDNA and PX-DNA, respectively, with a speed of 0.1 nm/ns. [C] PMF profile for a pulling velocity of 0.1 nm/ns using JE with red and sky blue colors indicating dsDNA and PX-DNA, respectively. [D, E] display the work profiles of DNase I dissociation from the binding region of dsDNA and PX-DNA with a speed of 1 nm/ns. [F] PMF profiles for dsDNA and PX-DNA in red and sky blue colors, respectively.

We hypothesize that SMD pulling trajectories impose restrictions on the DNase I’s rotation, indicating that SMD did not provide enough sampling to capture its rotational degrees of freedom, as also reported in the literature for DNA-protein interactions.^64^ This affects DNase I’s energetics, leading to an overestimation of absolute binding free energy Δ*G_b_*. To address this limitation, umbrella sampling was employed to improve sampling on DNase I’s rotational degrees of freedom, as discussed in the next section. Although JE overestimates the binding free energy values for dsDNA and PX-DNA, it correctly captures the binding affinity trend of DNase I, aligning with previous results.

### Calculation of Residence time using ***τ*** -RAMD

The short timescales of traditional unbiased molecular dynamics simulations make it difficult to estimate the residence time (*τ*) of DNA-DNase I interactions, which is essential for comprehending the biostability of DNA nanostructures. To overcome this, we used *τ* -Random Acceleration Molecular Dynamics (*τ*-RAMD), a method that is useful for calculating residence times and dissociation rates in protein-small molecule systems.^64–67^ *τ*-RAMD accelerates unbinding events by applying randomly oriented forces (shown in colored arrows in Figure 6A, B, and C) to the ligand’s (DNase I) center of mass (COM). The random force orientation ensured unbiased sampling of ligand unbinding pathways. For every 100 fs, the DNase I’s displacement was measured, and a new random force orientation was applied if the distance was less than 0.025 Å. The simulation was terminated upon reaching a DNA-DNase I COM separation of 50 Å. 15 RAMD dissociation trajectories were generated using the starting configurations from the final snapshots of the equilibrium unbiased simulations. A random force of magnitude 1500 kj.mol^-1^.nm^-2^ was applied to find the DNA-DNase I dissociation rates, and 10 ps intervals were used to record the snapshots during *τ*-RAMD. Based on the previous findings of structural and energetic properties, we conclude that the JX topoisomer’s binding affinity is intermediate between dsDNA and PX-DNA; thus, we applied the *τ*-RAMD only to the dsDNA and PX-DNA to calculate their residence times. Our findings from 15 RAMD trajectories indicate that the average DNase I residence time is 0.19 ns for PX-DNA and 0.62 ns for dsDNA, revealing a three-fold lower residence time for PX-DNA. Consequently, DNase I dissociates faster from PX-DNA backbones, indicating higher nuclease resistance (Figure 6). Figure S2 (SI) shows the cumulative and residence time distributions of the RAMD trajectories.

**Figure 6:**
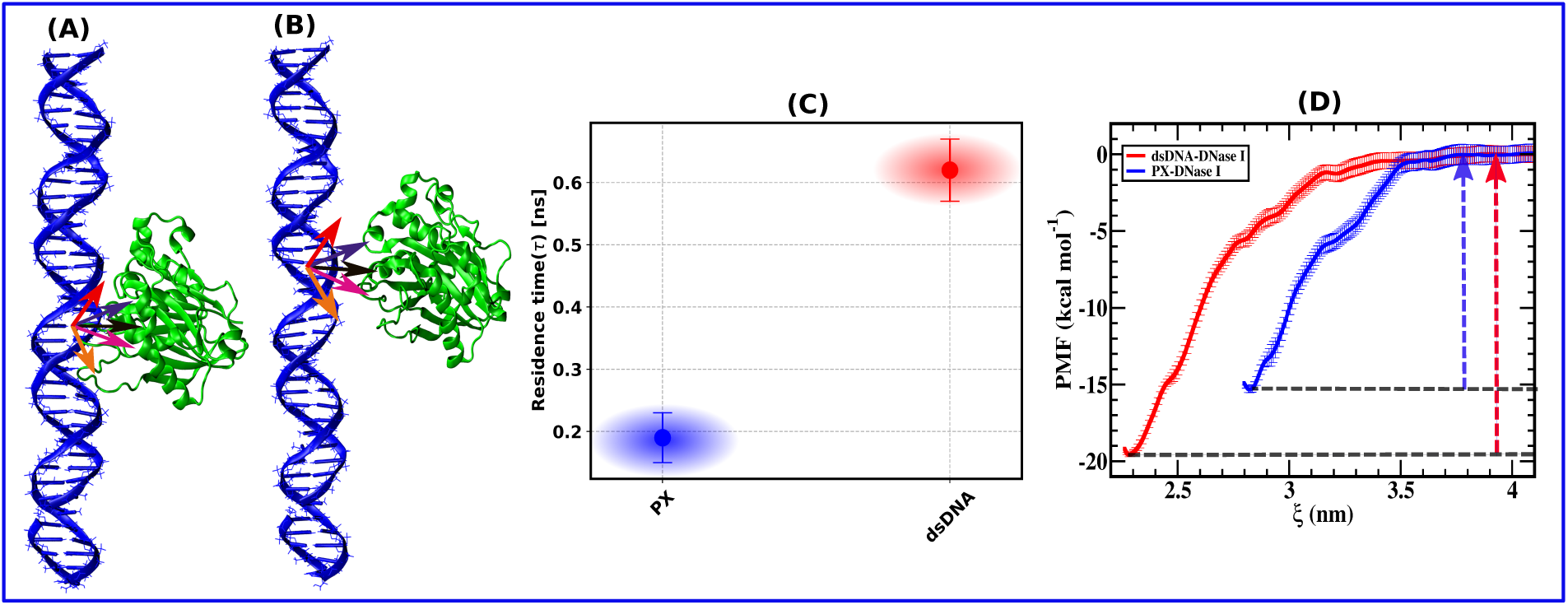
(A) Initial confirmation from the *τ*-RAMD simulation for dsDNA-DNase I complex, showing without water and ions for clarity. (B) Final confirmation of the *τ*-RAMD simulation for the dsDNA-DNase I complex. (C) The calculated residence time (*τ*) of DNase I dissociation from the binding regions of dsDNA and PX-DNA nanostructures, the DNase I’s center of mass (COM), was directed in an arbitrary direction. (D) PMF plot from the umbrella sampling (US) simulation of dsDNA-DNase I complex (in red color) and PX-DNase I complex (in blue color). The PMF is a function of the distance (*ξ*) between the center of mass of DNase I and the binding sites of the dsDNA and PX-DNA.

### Binding Free Energy Calculation using Umbrella Sampling

Umbrella Sampling (US) was used to obtain the binding free energy profile of DNase I with native dsDNA and PX-DNA, with the COM distance (*ξ*) between the DNase I and dsDNA/PX-DNA binding sites as the reaction coordinate. Steered MD simulations generated initial configurations for ∼ 40-50 US windows (with 0.5 Å spacing) using a 0.0001 nm/ps pulling rate. Each of the US windows was run for 20 ns to compute the PMF profile. To assess DNase I’s binding affinity to dsDNA and PX-DNA, we computed the PMF profile (Figure 6D). DNase I must overcome the free energy barrier to dissociate from the DNA-bound state, which is measured from the global minimum to the maximum PMF value. Binding free energies are 19.65 ± 0.49 kcal/mol for dsDNA and 15.41 ± 0.63 kcal/mol for PX-DNA, indicating a lower affinity for PX-DNA. The global energy minimum for the dsDNA is at 2.28 nm, compared to 2.82 nm for PX-DNA, indicating that dsDNA has stronger binding due to the shortest distance (deeper penetration in dsDNA grooves) and making it more susceptible to degradation under physiological conditions. The weaker DNase I binding to PX-DNA also correlates with a shorter residence time (*τ*), which is consistent with the Free Energy Landscape (FEL) results discussed in the following section.

### Free Energy Landscape (FEL)

Previous studies identified R41 and Y76 as the two key active-site residues essential for DNase I’s nuclease activity.^68^ These residues interact with DNA to cleave phosphodiester bonds. Therefore, calculating the minimum distances between R41/Y76 and DNA bases is crucial for assessing the binding affinity. We computed the free energy landscape (FEL) from these two distances for all the six DNA-DNase I complexes as shown in Figure 7(A-F) using equation-8 as follows-

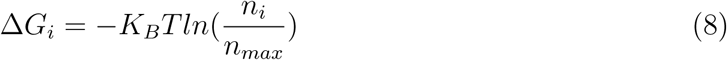

where *n_i_*denotes the population of the i^th^ bin, *n_max_* denotes the population of the most populated bin, and *K_B_*represents the Boltzmann constant. T is the temperature of each simulated system.

**Figure 7:**
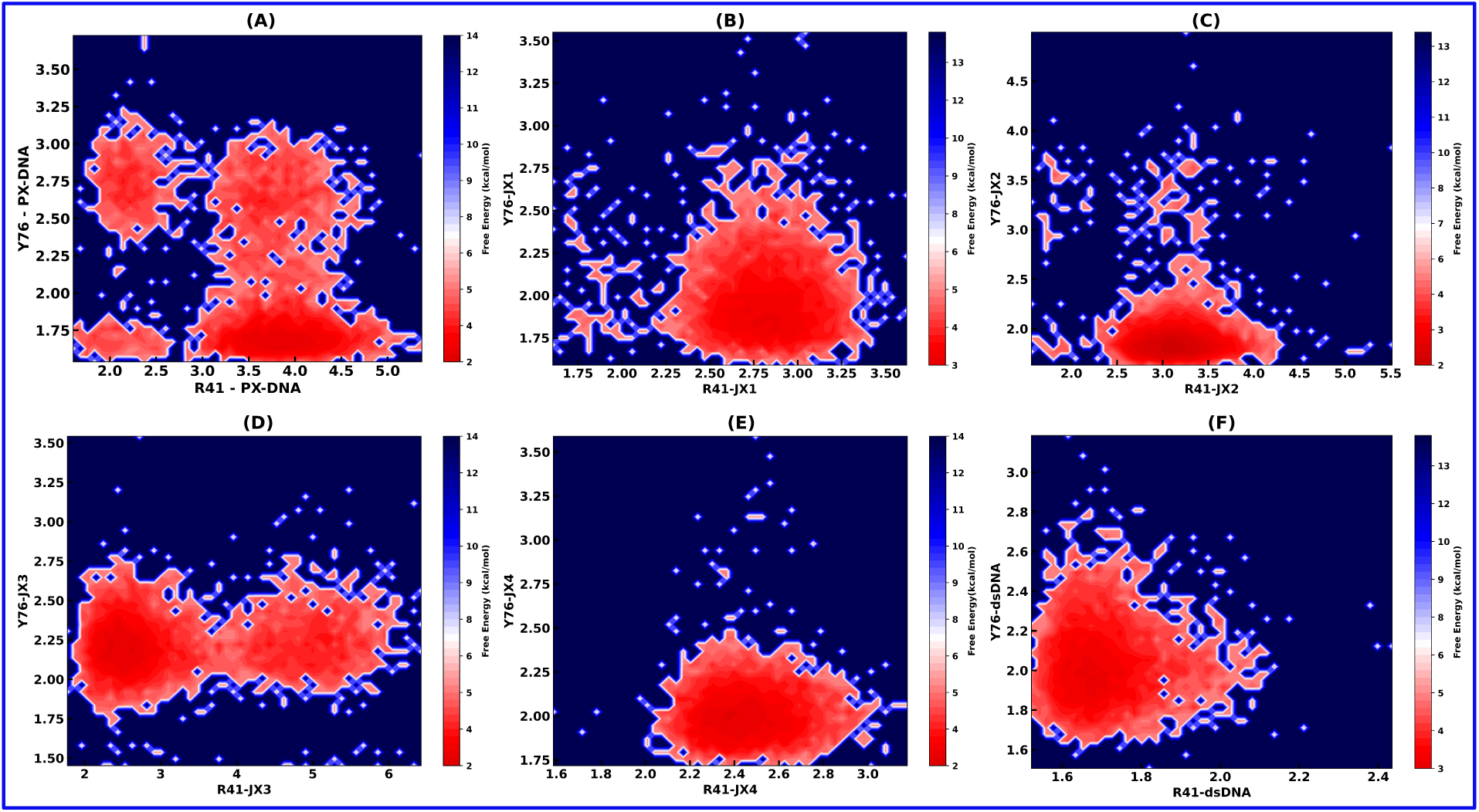
The free energy landscape of DNase I residues (R41 and Y76) and DNA nanostructure residues quantifies enzyme penetration into DNA grooves (x/y-axis in Å units). Panels (A-F) represent PX-DNA, JX1, JX2, JX3, JX4, and dsDNA, respectively, with the side color bar indicating free energy in kcal/mol unit.

The global energy minima in FEL plots (in deep red color) reveal that R41 and Y76 prefer longer distances (4.0 Å, 1.75 Å) in PX-DNA, reducing its degradability, whereas they maintain closer proximity in native dsDNA (1.7 Å, 2.0 Å), enhancing binding affinity. This suggests that DNase I can efficiently bind and cleave native dsDNA by penetrating its minor groove via strong electrostatic interaction and hydrogen bonding. JX(1,2,3 and 4))-DNA motifs exhibit intermediate stability, indicating that increased crossovers enhance resistance to DNase I nuclease. The distance-based FEL provides a better understanding of the mechanism behind these variations in binding affinities. A residue-residue contact map between DNA nanostructure and DNase I residues is shown in Fig.S3 (SI).

### Mechanical Properties

One of the key features of DNA structure is the twist angle, ^69^ with native dsDNA typically showing ∼10.5 base pairs per helical turn. We observe that the twist angle increases as the number of crossover points increases in PX/JX DNA motifs. Therefore, the native dsDNA has the lowest twist angle (∼32*^◦^*) with no crossovers, while PX-DNA has the highest twist angle (∼36*^◦^*) with six crossovers as listed in Table-3. JX topoisomers exhibit intermediate values of twist in the order PX*>*JX4*>*JX3*>*JX2*>*JX1*>*dsDNA.

**Table 3:**
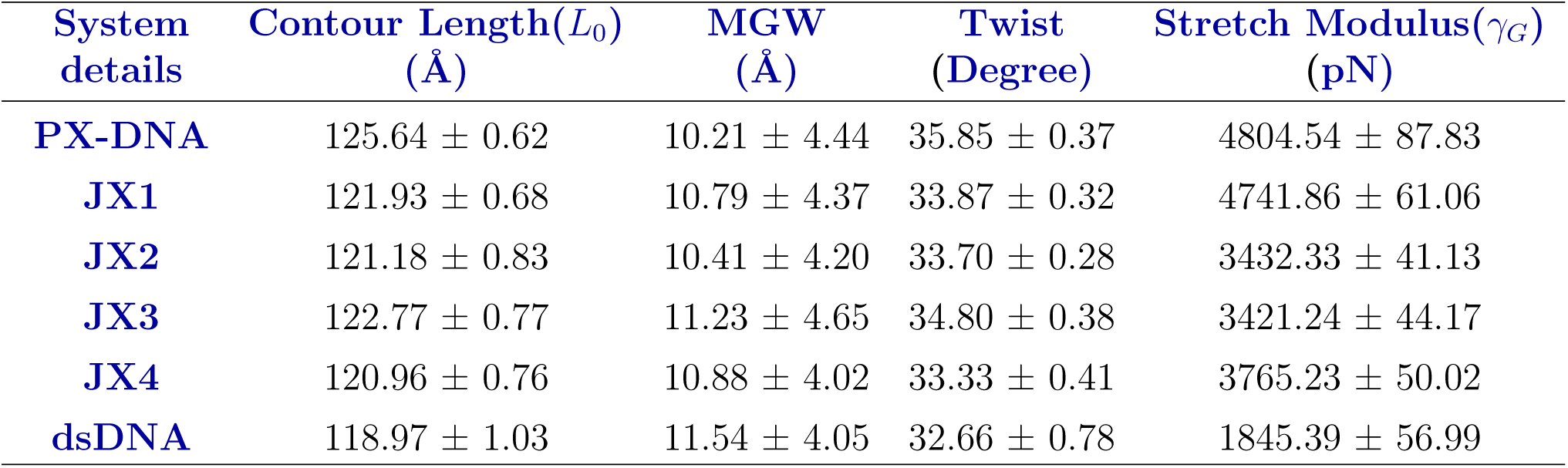
Mechanical Properties of native dsDNA and PX/JX-DNA nanostructures using contour length distribution from the last 200 ns all-atom simulation trajectory at 300 K and the Helical parameters (MGW and Twist).

| System details | Contour Length($L_0$)<br>(Å) | MGW<br>(Å) | Twist<br>(Degree) | Stretch Modulus($\gamma_G$)<br>(pN) |
| --- | --- | --- | --- | --- |
| **PX-DNA** | $125.64 \pm 0.62$ | $10.21 \pm 4.44$ | $35.85 \pm 0.37$ | $4804.54 \pm 87.83$ |
| **JX1** | $121.93 \pm 0.68$ | $10.79 \pm 4.37$ | $33.87 \pm 0.32$ | $4741.86 \pm 61.06$ |
| **JX2** | $121.18 \pm 0.83$ | $10.41 \pm 4.20$ | $33.70 \pm 0.28$ | $3432.33 \pm 41.13$ |
| **JX3** | $122.77 \pm 0.77$ | $11.23 \pm 4.65$ | $34.80 \pm 0.38$ | $3421.24 \pm 44.17$ |
| **JX4** | $120.96 \pm 0.76$ | $10.88 \pm 4.02$ | $33.33 \pm 0.41$ | $3765.23 \pm 50.02$ |
| **dsDNA** | $118.97 \pm 1.03$ | $11.54 \pm 4.05$ | $32.66 \pm 0.78$ | $1845.39 \pm 56.99$ |

Another crucial factor in DNase I binding or nuclease resistance is the minor groove width (MGW).^70^ For instance, previous studies have shown that the minor groove width can be used to predict the rate at which DNase I cleaves DNA phosphodiester bonds.^70–73^ Native dsDNA has the widest MGW, followed by JX and PX-DNA motifs, with MGW decreasing as the crossover points increase, as listed in Table-3. A strong negative correlation exists between MGW and twist angle, with both factors directly linked to the number of crossover points in DNA nanostructures. Therefore, DNase I nuclease binds more readily to dsDNA due to its lower twist, wider MGW, and closer proximity to active residues. In contrast, narrower grooves in PX/JX-DNA motifs hinder DNase I binding, leading to weaker interactions and reduced cleavage efficiency.

### Stretch Modulus of DNA nanostructures from Contour Length Distributions

We calculated the DNA stretch modulus using a microscopic elastic rod model. The instantaneous contour length L, fluctuates around its equilibrium length *L*_0_, due to thermal energy. It turns out that the instantaneous contour length L is the sum of rises (center of mass distance between consecutive base pairs). The linear elastic rod model assumes that instantaneous contour length (L) fluctuations will give rise to restoring force. The restoring force (F) is proportional to (*L* − *L*_0_) and expressed as *F* (*L*) = −*γ*(*L* − *L*_0_)*/L*_0_, where *γ* is the rod’s stretch modulus.

Integrating the force with respect to L yields *E*(*L*) = *γ*(*L* − *L*_0_)^2^*/*2*L*_0_. If *l_i_* is the center-to-center distance between two consecutive base pairs, the average contour length is defined as 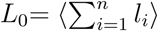 where ⟨⟩ represents the time average over all frames. Therefore, the contour length probability distribution can be expressed as a Gaussian as follows:

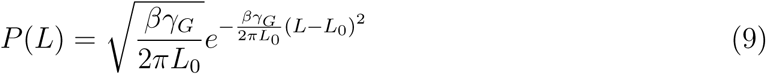

By taking the logarithm, equation-9 can be written as:

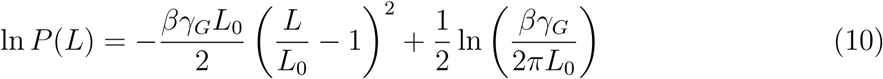

Where T is the equilibrated system’s temperature, *K_B_* is the Boltzmann constant, *L*_0_ is the time-averaged contour length, *γ_G_* is the stretch modulus, and 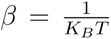. From the slope of the equation-10, we can determine the stretch modulus *γ_G_* of the dsDNA molecules.^74,75^ Using the above approach, we calculated the stretch modulus for native dsDNA and PX/JX-DNA nanostructures. The contour length of the PX/JX DNA was determined by summing the COM distances of consecutive slices along the long helical axis. The contour length distribution with simulation time for all the complexes is shown in Figure S4 in SI. Using equation-9,10 the stretch modulus was calculated, where PX-DNA exhibits the highest stretch modulus (*γ_G_*) of 4805 ± 87 pN, while dsDNA has the lowest *γ_G_* value of 1845.39 ± 56.9 pN at 150 mM salt concentration as listed in Table-3. These values align with previous studies ^76^ in bulk water in the absence of DNase I nuclease, which showed PX/JX-DNA stretch modulus between 1300-2300 pN in neutralized solutions. Maiti and coworkers have successfully used similar methodologies in their earlier studies.^38,39,75–78^ Naskar et al. showed that increasing monovalent salt concentration enhances DNA’s structural stability and mechanical rigidity^75^ in absence of DNase I. Our results reflect that the presence of both physiological salt concentration and DNase I enzyme significantly affects the structural stability and mechanics of dsDNA and PX/JX-DNA nanostructures. The stretch modulus was derived from the slope of the logarithmic fit of contour length distributions using equations-9,10, and the data set for each of the six systems contour length probability distribution and logarithmic fit is shown in Figure 8(A–F) and Figure S5(A–F)(SI), respectively. The JX topoisomers show intermediate *γ_G_* values based on crossover points as shown in Figure 9 (values are listed in Table-3).

**Figure 8:**
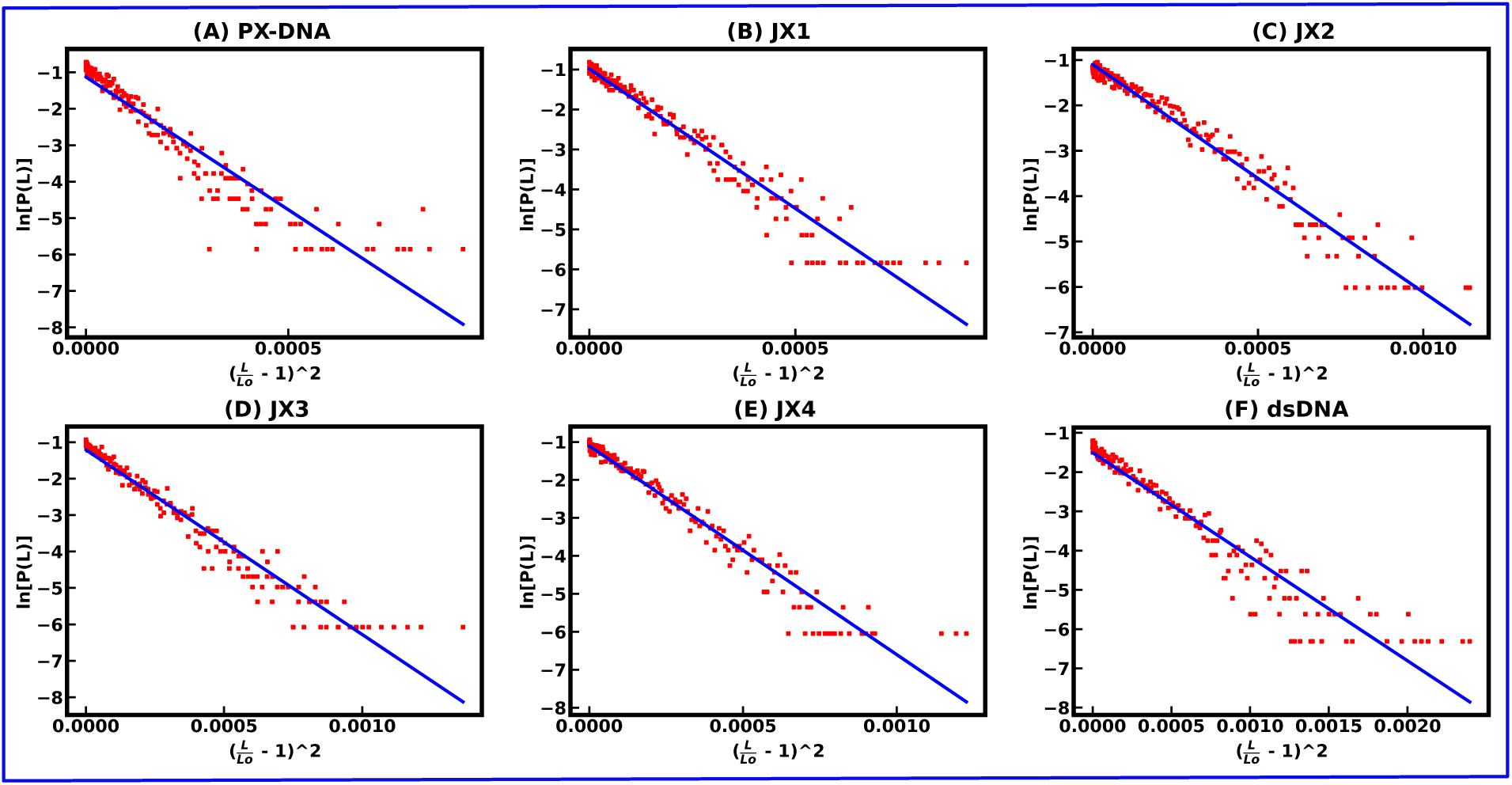
Representation of plots for the contour length fitting as ln *P* (*L*) vs 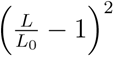 using the last 200 ns of the 300 ns long all-atom MD simulation trajectory for all six DNA-DNase I complexes at 300 K as displayed by (A-F) respectively.

**Figure 9:**
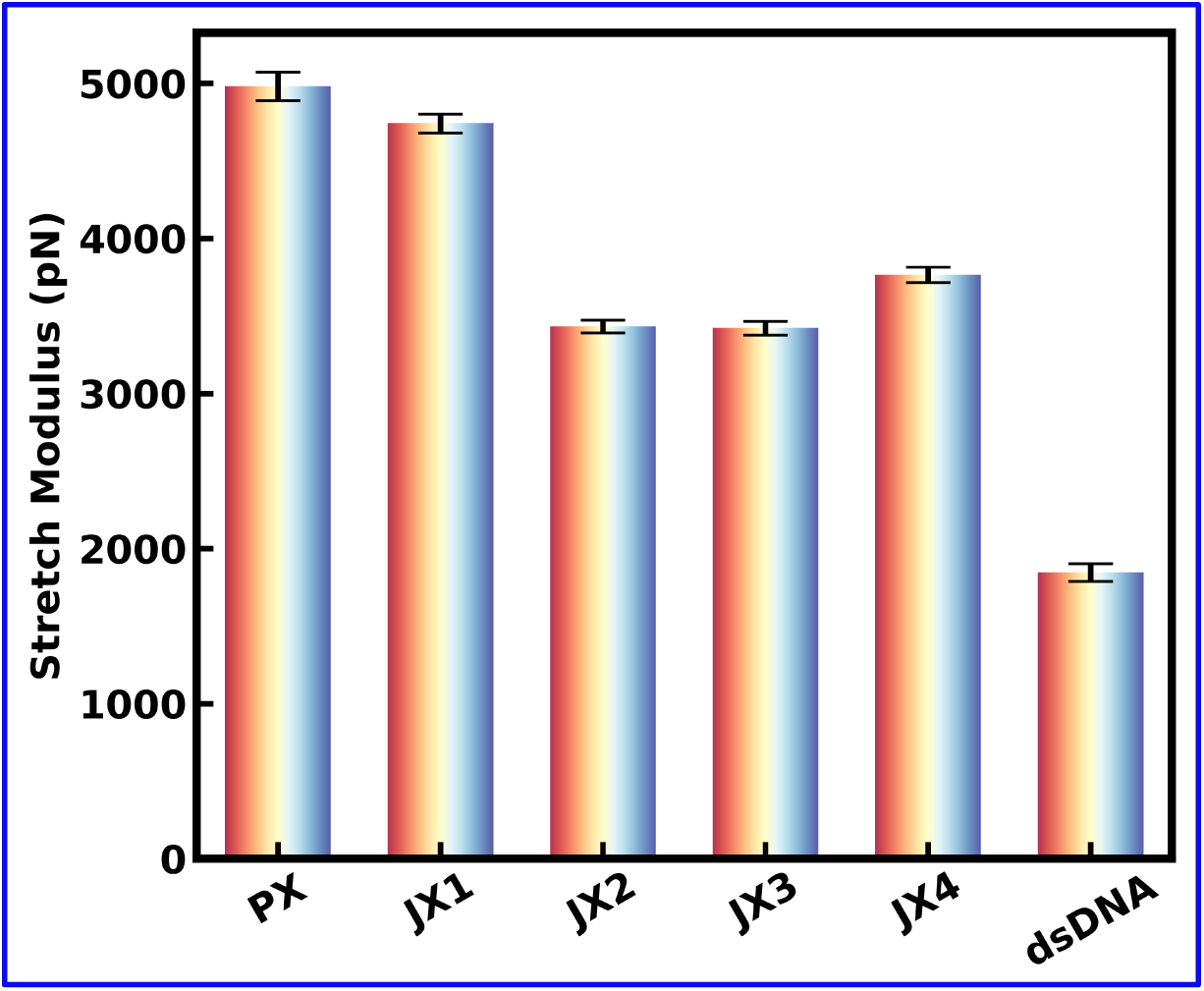
Stretch modulus of all six DNA nanostructures from their contour length distribution using the last 200 ns of the 300 ns long MD simulation trajectory, providing information on their flexibility when DNase I enzyme binds to the DNA nanostructures.

## Conclusion

In this work, we used atomistic MD simulations to reveal how DNase I binding affinity to the native dsDNA and PX/JX-DNA nanostructures is driven by key factors such as number of crossover points, helical twist, minor groove width (MGW), and stretch modulus (*γ_G_*), which influence nuclease resistance. Our findings highlight that DNase I activity on DNA is closely linked to the number of crossovers between neighboring helices, as well as the number of HBs and non-bonded interactions. Previous work^79,80^ has shown that DNase I needs a substrate that is normally 6–8 base pairs, but the double helical area between consecutive crossovers in PX-DNA is limited to 5 or 6 base pairs (alternating half-turns) because crossovers happen at every half-turn. These shorter binding regions of PX-DNA exhibit higher nuclease resistance than the native dsDNA and JX-DNA. This resistance is attributed to PX-DNA’s higher twist angle, stretch modulus, and narrower MGW. Thus, the shorter binding region and residence time indicate that DNase I cannot firmly grip or remain near PX-DNA residues for long.

The binding free energy (PMF) profiles from the JE and Umbrella Sampling (US) simulation confirm DNase I’s strong binding preference for native dsDNA, while PX-DNA’s weaker binding affinity correlates with its enhanced resistance to DNase I degradation. Our study reinforces the idea that crossover-dependent stability can be engineered into DNA nanostructures for improved biostability, increasing robustness for biological applications involving PX-DNA motifs.

Strategically placing crossovers in vulnerable regions can improve DNA nanostructure stability, resulting in ”tunable” biostability for controlled degradation, which is ideal for drug delivery. Further quantitative research is required to explore how DNA sequences influence DNase I binding dynamics and kinetics. Overall, this work underscores the importance of incorporating crossover points, MGW, stretch modulus, and helical twist in the design of DNA nanostructures. By considering these factors, the goal is to improve applications in biomedicine and other fields of DNA nanotechnology while also increasing the design’s purpose.

## Computational Methodology

### MD Simulation Details

The initial structure of the native dsDNA was built using NAB code, and the initial coordinates of PX/JX-DNA structures were obtained from previous work by Maiti et al. ^39^ The structure of the DNase I enzyme was obtained from the Protein Data Bank (PDB ID: 1DNK).^80^ The DNA-DNase I complex structures were then solvated in a cubic box with a buffer length of 15 Å using the TIP3P^81,82^ water model with xLEAP module of AMBER20.^56,83,84^ In this way, all atoms of the solute molecules will be 15 Å apart from the edge of the water box. The negative charges of the phosphate backbones were neutralized by adding the appropriate number of Na^+^ ions. To achieve systems with the desired salt concentration of 0.15 M, we added appropriate numbers of Na^+^ and Cl*^−^* molecules. Periodic boundary conditions were used in all three dimensions to mimic the bulk properties of the system. All MD simulations were carried out using PMEMD.cuda module of AMBER20. The interactions of DNA-DNase I complex atoms and TIP3P waters were defined by leaprc.DNA.bsc1 forcefield.^85^ For the monovalent ions (Na^+^, Cl*^−^*), the Joung-Chetham ion parameters were chosen. ^86^ Following that, the energy of the solvated systems was minimized using the steepest descent method for the first 4000 steps, followed by the conjugate gradient method for the next 6000 steps. All of the solute atoms were held fixed by a harmonic potential with a force constant of 500 kcal.mol^-1^.Å^-2^ so that ions and water molecules can redistribute themselves from their initial ordered configuration to remove any bad contacts. The systems were gradually heated in four steps: first, from 10 K to 50 K for 6000 steps, then, from 50 K to 100 K for 12,000 steps, then from 100 K to 200 K for 10,000 steps, and finally, from 200 K to 300 K for 10,000 steps. Positional restraints were applied to the heavy atoms of the solute during heating with a force constant of 50 kcal.mol^-1^. Å^-2^. Following heating, the systems underwent a 5 ns NPT simulation with a Berendsen weak coupling method^87,88^ to adjust the box size with a target pressure of 1 bar. All bonds involving hydrogens were constrained using the Shake algorithm.^89^ This allowed us to use an integration time step of 2 fs. The electrostatic interactions were treated using the Particle Mesh Ewald method^90^ with a cutoff of 10 Å. Following the equilibration, the systems were simulated using a Langevin thermostat^88,91^ with coupling constant 1 ps at 300 K for 300 ns simulation time. Similar simulation methodologies at ambient temperatures have been implemented in some of the previous studies.^78,92–96^

Simulation trajectories were analyzed using VMD^97^ for 3D visualization and CPPTRAJ^55^ for modeling and analysis. Data visualization was done with Grace, PyMOL, and Matplotlib, while functional fittings used Gnuplot and Grace.

### Umbrella Sampling Simulation Setup

The final structures of the DNase I bound to dsDNA and PX-DNA complexes were obtained from the 300 ns unbiased all-atom MD simulations. Water and ion (Na^+^ and Cl*^−^*) molecules were removed, leaving only the DNase I attached to the DNA strands of dsDNA and PX-DNA. The systems were solvated using a bigger water box of dimension 120 × 65 × 130 Å^3^ along x, y, and z directions, respectively (the x-axis is the direction of pulling). It was then initially neutralized with counterions, and then more Na^+^ and Cl*^−^* ions were added to get the desired salt concentration of 0.15 M. A similar process described in the previous section was used to minimize energy, followed by 10 ns NPT MD simulations to achieve equilibrium at 300 K.

The DNase I was then forced to dissociate from the equilibration phase by moving its center of mass (COM) along the x-axis to produce initial configurations for each umbrella sampling window using the quasi-equilibrium steered molecular dynamics (SMD). In SMD, the DNase I was moved slowly at a speed of 0.0001 nm/ps, with the help of a spring force with a force constant of 3000 kJ.mol*^−^*^1^.nm*^−^*^2^. During SMD pulling, the DNase I was moved to 25 Å away from the DNA binding sites in steps of 0.5 Å. To hinder the DNA from being pulled along with the DNase I during the pulling process, the heavier atoms of the DNA were restrained by harmonic restoring forces, which had a force constant of 1000 kJ.mol.*^−^*^1^.nm*^−^*^2^. Constraints were imposed to shift the enzyme away from the DNA binding sites in a direction perpendicular to the long axis of the DNA helix, hindering it from rotating around the long axis of DNA. Using umbrella sampling to improve the sampling of the energetically unfavorable regions enabled us to compute the potential of mean force (PMF) by applying an external harmonic bias potential. In this study, the PMF profiles were generated using the GROMACS with the WHAM code.^98^

## Data availability

The data validating the findings of our simulation study are available upon request to the corresponding author.

## Acknowledgement

S.M. acknowledges the SRF fellowship from CSIR, India. P.K.M. thanks SERB, India, for financial through CRG/2021/003659. A.R.C. acknowledges support from the National Institutes of Health (NIH) through the National Institute of General Medical Sciences (NIGMS) under award number R35GM150672. We are thankful to Prof. Himanshu Joshi for the valuable suggestions.

## TOC Graphic

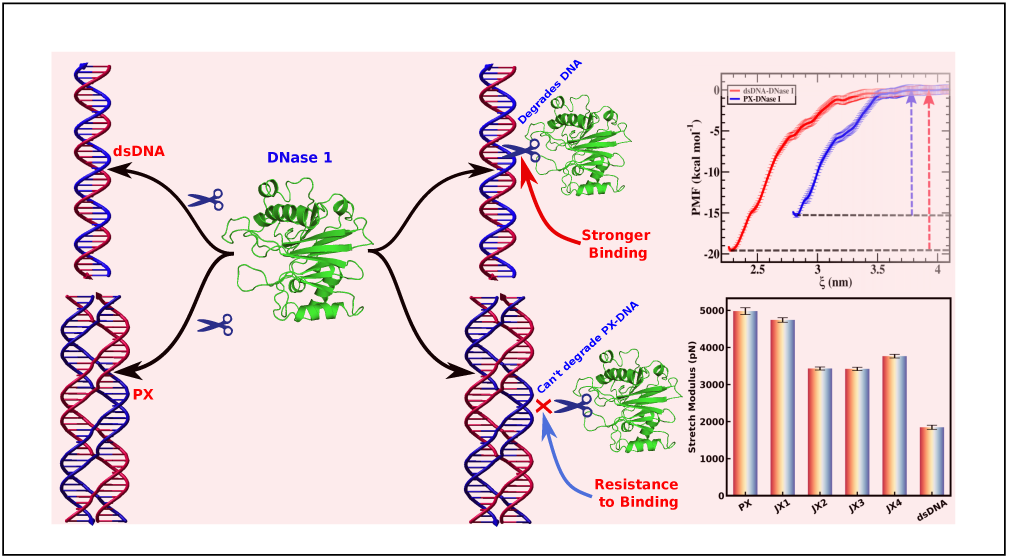

